# The Anti-histamine Azelastine, Identified by Computational Drug Repurposing, Inhibits SARS-CoV-2 Infection in Reconstituted Human Nasal Tissue In Vitro

**DOI:** 10.1101/2020.09.15.296228

**Authors:** Robert Konrat, Henrietta Papp, Valéria Szijártó, Tanja Gesell, Gábor Nagy, Mónika Madai, Safia Zeghbib, Anett Kuczmog, Zsófia Lanszki, Zsuzsanna Helyes, Gábor Kemenesi, Ferenc Jakab, Eszter Nagy

## Abstract

**Background:** The COVID-19 pandemic is an enormous threat for healthcare systems and economies worldwide that urgently demands effective preventive and therapeutic strategies. Unlike the development of vaccines and new drugs specifically targeting SARS-CoV-2, repurposing of approved or clinically tested drugs can provide an immediate solution.

**Methods:** We applied a novel computational approach to search among approved and clinically tested drugs from the DrugBank database. Candidates were selected based on Shannon entropy homology and predefined activity profiles of three small molecules with proven anti-SARS-CoV activity and a published data set. Antiviral activity of a predicted drug, azelastine, was tested *in vitro* in SARS-CoV-2 infection assays with Vero E6 monkey kidney epithelial cells and reconstituted human nasal tissue. The effect on viral replication was assessed by quantification of viral genomes by droplet digital PCR.

**Findings:** The computational approach with four independent queries identified major drug families, most often and in overlapping fashion anti-infective, anti-inflammatory, anti-hypertensive, anti-histamine and neuroactive drugs. Azelastine, an histamine 1 receptor-blocker, was predicted in multiple screens, and based on its attractive safety profile and availability in nasal formulation, was selected for experimental testing. Azelastine significantly reduced cytopathic effect and SARS-CoV-2 infection of Vero E6 cells with an EC_50_ of ∼6 μM both in a preventive and treatment setting. Furthermore, azelastine in a commercially available nasal spray tested at 5-fold dilution was highly potent in inhibiting viral propagation in SARS-CoV-2 infected reconstituted human nasal tissue.

**Interpretations:** Azelastine, an anti-histamine, available in nasal sprays developed against allergic rhinitis may be considered as a topical prevention or treatment of nasal colonization with SARS-CoV-2. As such, it could be useful in reducing viral spread and prophylaxis of COVID-19. Ultimately, its potential benefit should be proven in clinical studies.

**Funding:** provided by the Hungarian government to the National Laboratory of Virology and by CEBINA GmbH.

## INTRODUCTION

The COVID-19 (Coronavirus Disease 2019) pandemic represents a worldwide threat to public health and demands immediate response to rapidly develop effective preventive and therapeutic approaches. Given the immense time pressure, the established pathway to drug discovery and development is not feasible, and repurposing of clinically approved drugs is an attractive alternative.

Drug repurposing/repositioning means identifying new indications for existing drugs. It is a promising option for rapid identification of new therapeutic agents.^1^ There are three main approaches: 1., rational selection based on the knowledge of mode-of-action of a given drug and the pathogenesis of the disease, 2., large-scale high throughput screening in *in vitro* assays for the required function, and 3., computational prediction using drug data bases. All of these have been applied to COVID-19 and several drugs are already being tested in clinical studies. The only drug that has achieved significant survival benefit against COVID-19 so far is dexamethasone.^2^ However, it is not a real drug repurposing example, since the original indication and known anti-inflammatory mode of action of glucocorticoids has been utilized to tame the overshooting inflammation that is characteristic of COVID-19. Several other anti-inflammatory drugs are being tested clinically, e.g. monoclonal antibodies against pro-inflammatory cytokines, such as IL-6. A rational approach, a kind of semi-drug repurposing case is the use of directly antiviral compounds, such as remdesivir. Remdesivir was an experimental stage (Phase 3) drug at the onset of the COVID-19 pandemic, and originally developed against the Ebola virus, which is also an RNA virus like SARS-CoV-2 and has proven potent activity against SARS-CoV-2 both in *in vitro* and *in vivo* infection models (reviewed in ^3^). Although considered as a therapy in COVID-19, its clinical benefit is modest.^3^ A classic example of true drug repurposing against COVID-19 is chloroquine/hydroxychloroquine, anti-malaria drugs that were shown to have activity against coronaviruses from previous pandemics and also against SARS-CoV-2.^4^ In addition, hydroxychloroquine is also used as an anti-inflammatory drug in autoimmune patients.^5^ Besides the concern about potential severe side-effects, well-designed sufficiently powered clinical studies proved hydroxychloroquine ineffective in preventing or treating COVID-19 in patient populations with different disease severity.^6^ The discrepancy between the *in vitro* activity and the clinical efficacy of these drugs might be explained by recent data demonstrating the lack of potency against SARS-CoV-2 infection in respiratory epithelial cells.^7,8^ Another rational approach, but truly drug repurposing cases are the angiotensin converting enzyme 2 (ACE2) inhibitors and angiotensin receptor antagonists widely used in anti-hypertensive therapy. Since ACE2 is the receptor for SARS-CoV-2 entry, but also shown to be protective against lung damage in animal models ^9^, both beneficial and potential disease enhancing effects are considered.

While not purposefully designed, it is known that the high therapeutic efficacy of some of the oldest and most valuable drugs is due to their interactions with multiple targets causing synergistic beneficial effects.^10,11^ This pleiotropic activity profiles of known drugs were found serendipitously, or are even unknown. Modern drug discovery is gradually shifting its strategy and starting to embark on multi-target approaches.^12,13^ This is also the basis of large-scale high throughput drug screening and computational prediction programs, both already applied to COVID-19 and several drugs have been recently identified with anti-SARS-CoV-2 activity *in vitro*.^8,14,15^

Here we have applied a novel computational prediction approach relying on biochemical pathway-based strategy. Key to our strategy is the poly-pharmacological hypothesis, that drugs simultaneously interact and interfere with numerous targets and thereby rewire the biochemical pathway networks. The drug identification problem is thus defined by finding a drug that matches a pre-defined pathway modulation profile. The starting point of our approach is a Shannon entropy-based description of the inherent chemical features of small molecules (drugs) with proven activity against SARS-CoV-2 and implications for protein target specificity. With this unique descriptor, it is possible to identify hidden and unexpected drug similarities and identify novel protein targets involved in the underlying biochemical pathways.

This prediction approach has identified drug families and approved drugs, some with proven anti-SARS-CoV-2 activity or clinical efficacy. Our results highlight the role of anti-histamines as a promising anti-COVID-19 drug class. Most importantly, we provide evidence that the histamine 1 (H1) receptor blocker azelastine, widely used in allergic rhinitis therapy in a nasal spray formulation, is effective against SARS-CoV-2 infection.

## METHODS

### Computational method to identify putative COVID-19 drugs

The identification strategy described here is based on the hypothesis that drugs with similar biochemical pathway (activity) profiles will address similar disease areas. Our computational approach was based on the Shannon-Entropy Descriptor (SHED) concept in which the chemical structure is converted into a 2D topological graph where nodes correspond to the atoms in the drug and edges connecting two nodes indicate the existence of a chemical bond. The most important feature of the SHED approach is, that seemingly different molecules with significantly different 2D molecular structures can nevertheless have similar Shannon entropy vectors resulting in similar biochemical (pharmacological) activity patterns. Details of the computational approach are given in the Supplementary Material 1.

The starting point in our search strategy was the selection of a desirable pathway profile and the subsequent search for clinically approved drugs that match this predefined activity pattern (Fig. 1). Relevant experimental information was available from a recent analysis of the KEGG pathways involved in SARS-CoV-2 infection.^16^ Secondly, we employed pathway information we predicted for drugs shown to be active against SARS-CoV and/or SARS-CoV-2. Three experimentally verified and characterized compounds, hydroxychloroquine ^17^, SSAA09E2 {N-[[4-(4-methylpiperazin-1-yl)phenyl]methyl]-1,2-oxazole-5-carboxamide} and SSAA09E3 {N-(9,10-dioxo-9,10-dihydroanthracen-2-yl)benzamide} ^18^ were employed. Hydroxychloroquine reduces endosomal acidification, SSAA09E2 acts by blocking early interactions of SARS-CoV with its receptor, the angiotensin converting enzyme 2 (ACE2), shared by SARS-CoV-2 and SSAA09E3 prevents fusion of the viral membrane with the host cellular membrane. For all three selected ligands, the pathway profiles were calculated, and the 50 highest scoring pathways were considered for the analysis.

**Figure 1.**
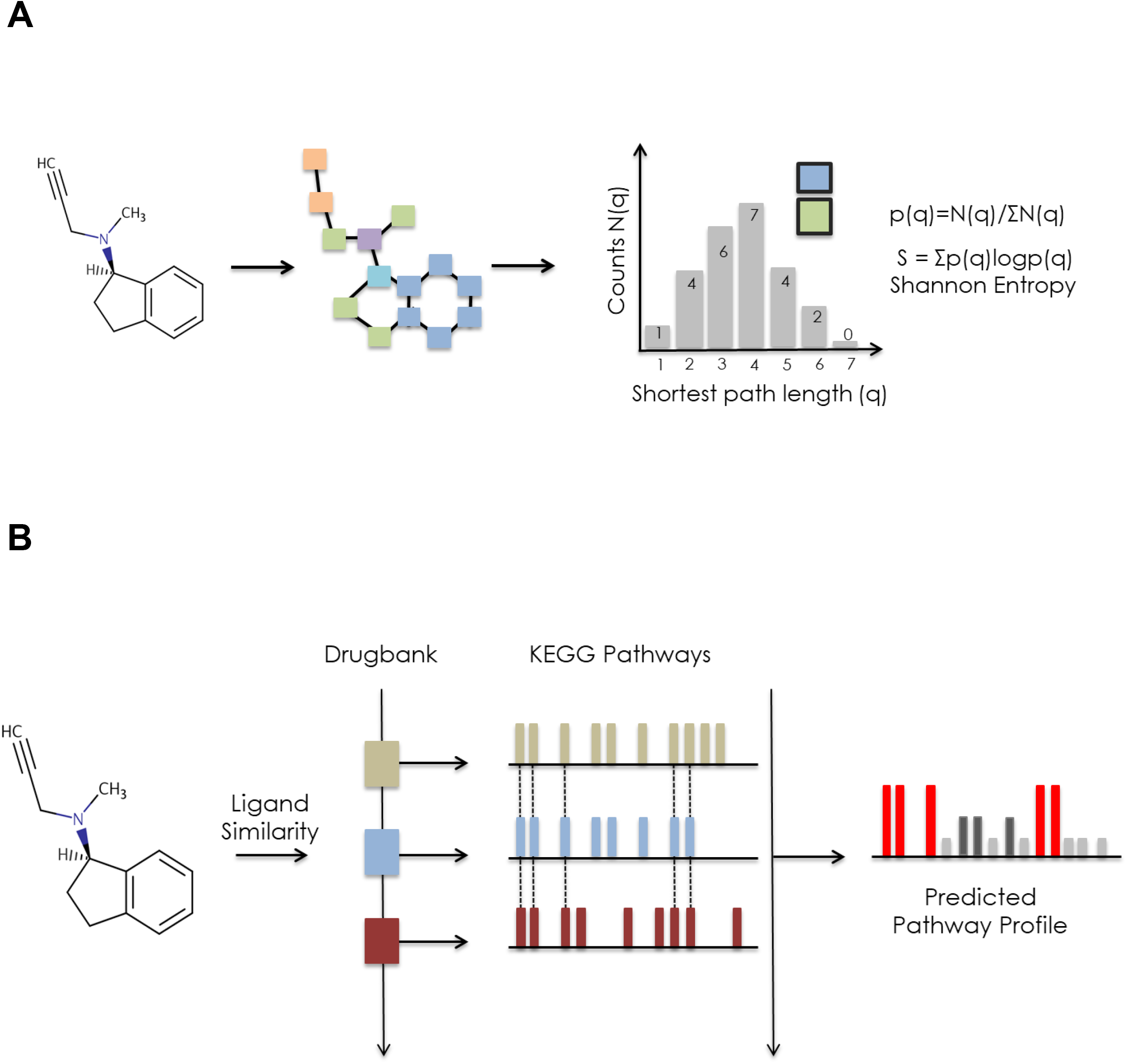
Schematic outline of drug prediction approach. **A:** Illustration of the first step: Shannon entropy approach for the description of small molecules. The 2D molecular structure is converted into a topological graph (network) in which every atom (node) is attributed with an atom type (feature). The shortest path length is calculated for each and every atom pair (feature pair). The shortest path length distribution N(q) of a given feature pair (i) is converted into a probability p(q) from which the Shannon entropy Si is calculated. The Shannon entropy descriptor SHED of a molecule is therefore the Shannon entropy vector S vector comprising entropy values for the individual feature pairs. **B:** Illustration of the second step: SHED-based prediction of pathway profiles. The Shannon entropy vector of a molecule is used to screen the DrugBank database for SHED analogs. DrugBank analogs are ranked according to pairwise SHED similarities with the query molecule. Protein targets and the corresponding KEGG pathways of individual DrugBank entries are used to predict the pathway profile of the query molecule (most prevalent/relevant pathways are indicated in red).

### SARS-CoV-2 infection assay with VERO E6 cell line

Vero E6 (African green monkey kidney epithelial) cells (ATCC CRL-1586) were seeded on 96-well plates at 4.5 10^4^/well and used at approximately 90% confluence. Azelastine-HCl (SeleckChem, S2552) was used at concentrations ranging from 0·4 to 25 μM. In a preventive setting, immediately after adding azelastine in the culture medium, the cells were infected with SARS-CoV-2 (hCoV-19/Hungary/SRC_isolate_2/2020, Accession ID: EPI_ISL_483637) at a MOI of 0·01. After 30 min incubation in a humidified atmosphere of 5% CO_2_ at 37°C, the culture medium was removed and replaced with fresh culture medium containing azelastine at given concentrations. In the post-infection treatment setting, cells were infected for 30 min without azelastine, then the culture medium containing the virus was removed, and replaced with fresh medium with azelastine. 48 hours post infection the cytopathogenic effect was evaluated by microscopic observation and the supernatants were collected for viral RNA copy number quantification. The cytopathic effect was assessed semi-quantitatively based on Cytopathic Scores (CPS) ranging from 0 to 4; 0: no cytopathic effect, comparable to uninfected control, 4: CPE is as strong as in the infected control. Cytopathic effect was based on the appearance of “holes” in the otherwise confluent, homogenous layer of cells indicating cell death. Cell viability in the presence of the drugs (without infection) was determined based on cellular ATP content using the CellTiter-Glo® Luminescent Cell Viability Assay (Promega, cat#: G7572) according to the manufacturer’s instruction.

### Testing the anti-SARS-CoV-2 activity of azelastine containing nasal spray with reconstituted human nasal tissue

MucilAir™ human nasal tissue generated from healthy donors (Epithelix, Cat#: EP02MP) was infected with SARS-CoV-2 at MOI of 0·01 on the apical side. After 20 min incubation at 37°C in 5% CO_2_, the virus containing media was removed completely. A 5-times diluted (in MucilAir™ culture medium) solution of the Allergodil nasal spray (0·1% azelastine, Mylan) was added onto the apical side (200μl) for 20 min. Following the treatment, the diluted nasal spray was fully removed from the surface of the cells to provide a liquid-air interface and incubated for 24 hours. The 20-min treatment with the diluted Allergodil was repeated at 24 and 48 hours post infection (hpi). After 24, 48 and 72 hpi, the apical sides of the cells were washed for 15 min with MucilAir™ Culture medium, and the solution was collected to quantify the viral RNA copy number. The cells were also inspected under an inverted microscope at 48 and 72 hpi.

### Virus quantification with Droplet digitalPCR analysis

Total RNA was extracted from 100 μl culture supernatants or apical washes using Monarch^®^ Total RNA Miniprep Kit (Promega, Cat#: T2010S). For viral copy number quantification droplet digital PCR technology was applied (Bio-Rad Laboratories Inc. QX200 Droplet Digital PCR System). The primers and probes used were specific for the SARS-CoV-2 RdRp gene (Reverse primer: CARATGTTAAASACACTATTAGCATA, Forward primer: GTGARATGGTCATGTGTGGCGG, Probe: FAM-CAGGTGGAACCTCATCAGGAGATGC-BBQ). For all concentrations at least three replicates were prepared and the EC_50_ values were determined with nonlinear regression (log(inhibitor) vs. response – variable slope (four parameter)) using GraphPad Prism 8.4.3 (GraphPad Software, San Diego, California USA).

Experiments with the infective SARS-CoV-2 were performed in the BSL4 facility of the Szentágothai Research Centre, University of Pécs, Hungary, according to institutional regulations.

## RESULTS

### Definition of query pathway profiles for drug repurposing

We found a significant overlap among the three predicted and the experimental KEGG pathways (Supplementary Material 2 and Fig. 2A). The most prominent was the overlap between the SSAA09E2 and Hydroxychloroquine pathways (34 of the 50 best hits). 9 out of 50 pathways were depicted in all three prediction sets, and four of these were also described in the experimental data set published by Zhou et al.^16^ Altogether 12 of the 95 unique pathways in the three predicted sets were described experimentally as well (Fig. 2, Supplementary Material 2). The most commonly identified disease areas associated with the predicted pathways were infectious diseases: viral infections (e.g. measles, Hepatitis C, EBV, influenza, HSV), parasitic infections (e.g. Chagas, Leishmaniasis, Trypanosomiasis, malaria), bacterial infections (e.g. Tuberculosis, Salmonella, Shigella, Pertussis, ETEC) or cellular processes involved during infection (e.g. endocytosis, immune cell signaling) (Supplementary Material 2).

**Figure 2.**
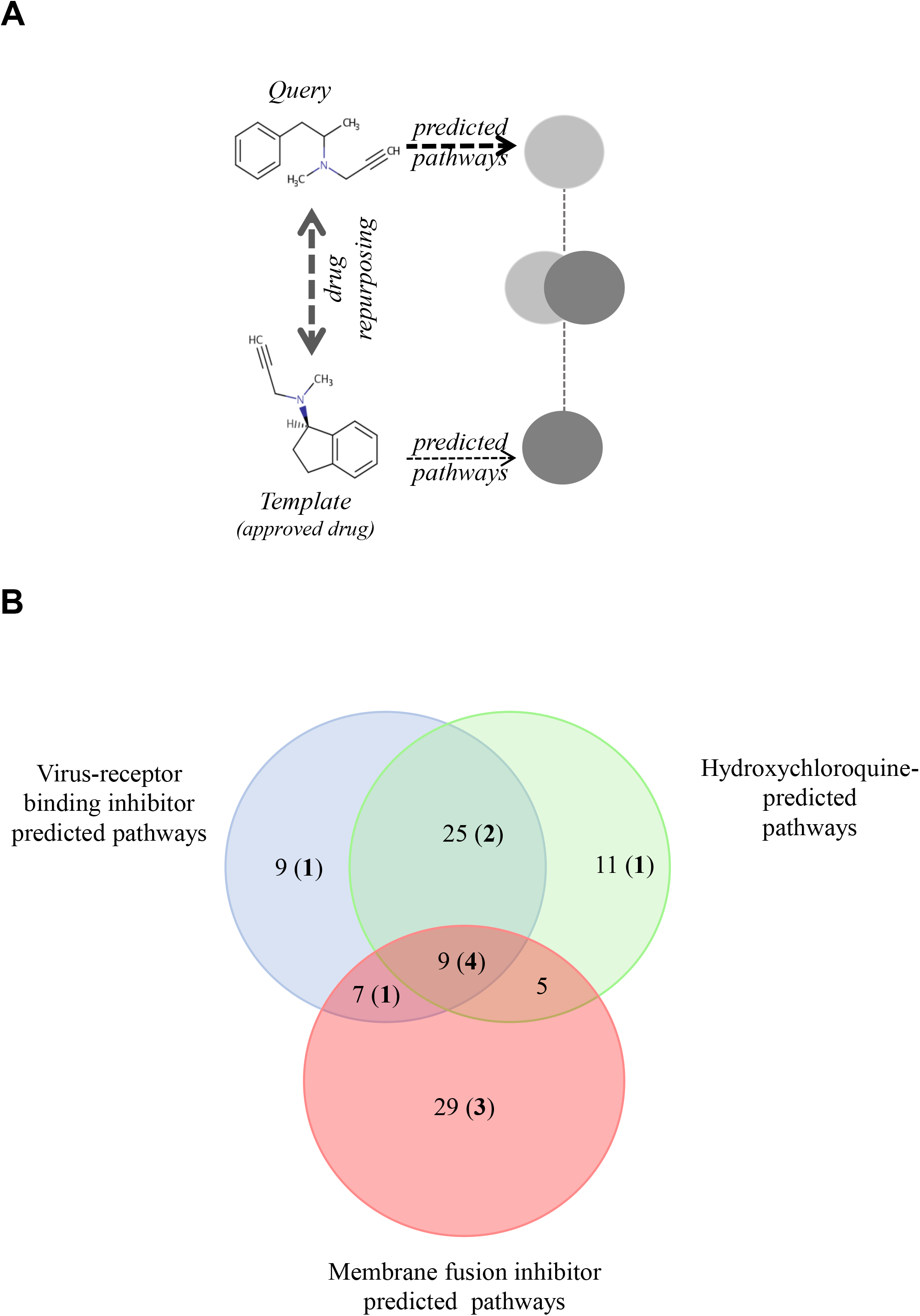
Pathway-based drug repurposing and overlap among three predicted and an experimentally verified pathway data set. (A) Illustration of pathway-based drug repurposing. A predefined pathway profile obtained for a query drug (outlined in Figure 1) is screened against a data set of template drugs with pre-calculated pathway profiles (for example, clinically approved drugs). Interesting drugs relevant for repurposing applications are obtained via maximizing pathway overlap. (B) Pathways predicted to be involved in SARS-CoV-2 infection were identified using three drugs shown to be active against SARS-CoV: a virus-receptor binding inhibitor: SSAA09E2, viral and cellular membrane fusion inhibitor: SSAA09E3 and hydroxychloroquine. Numbers in bold and parenthesis indicate the number of pathways also detected in an experimental data set by Zhou et al.^16^ KEGG pathways involved in this analysis are shown in Supplementary Material 2.

### Screening drugs for matching pathways

Next, a set of 2700 drugs - clinically tested, mostly approved and commercially available via SELLECKCHEM - was screened for matching pathways. Identified hits were stored as an ordered list and ranked based on the number of shared pathways (screened with the 50 best scoring pathways). The 100 top scoring drugs, predicted in four independent screens using the virus-receptor binding inhibitor SSAA09E2 (A), hydoxychloroquine (B), the membrane fusion inhibitor SSAA09E3 (C) and the experimental defined pathways for SARS-CoV-2 ^16^ (D) were considered for further analysis and are given in Supplementary Material 3 (listing the drugs identified in at least 2 screens and in major drug categories). The overlap between the predicted sets of drugs is in alignment with the corresponding predicted pathways. Approximately half of the drugs predicted based on the pathways of SSAA09E2 or hydoxychloroquine (that show approximately 2/3 overlap) were identical (Fig. 3). The main drug classes were anti-infectives (antivirals, antibiotics, antifungals), anti-histamines, anti-inflammatory drugs (non-steroids), anti-hypertensive drugs, diuretics (potassium channel blockers), analgesics and neuroactive drugs (mainly anti-psychotics) (Supplementary Material 3). Approximately 30% of the drugs identified both in data sets A and B were shared with those derived from the pathways from the experimental SARS-CoV-2 data set (D) (21 of 74, Fig. 3). We also found a significant drug overlap (approximately 30%) between data sets obtained for the viral fusion inhibitor SSAA09E3 and the experimental SARS-CoV-2 pathways. Here, the overlapping drugs were dominated by steroids of all kinds (anti-inflammatory glucocorticoids, progesterone-analogues).

**Figure 3.**
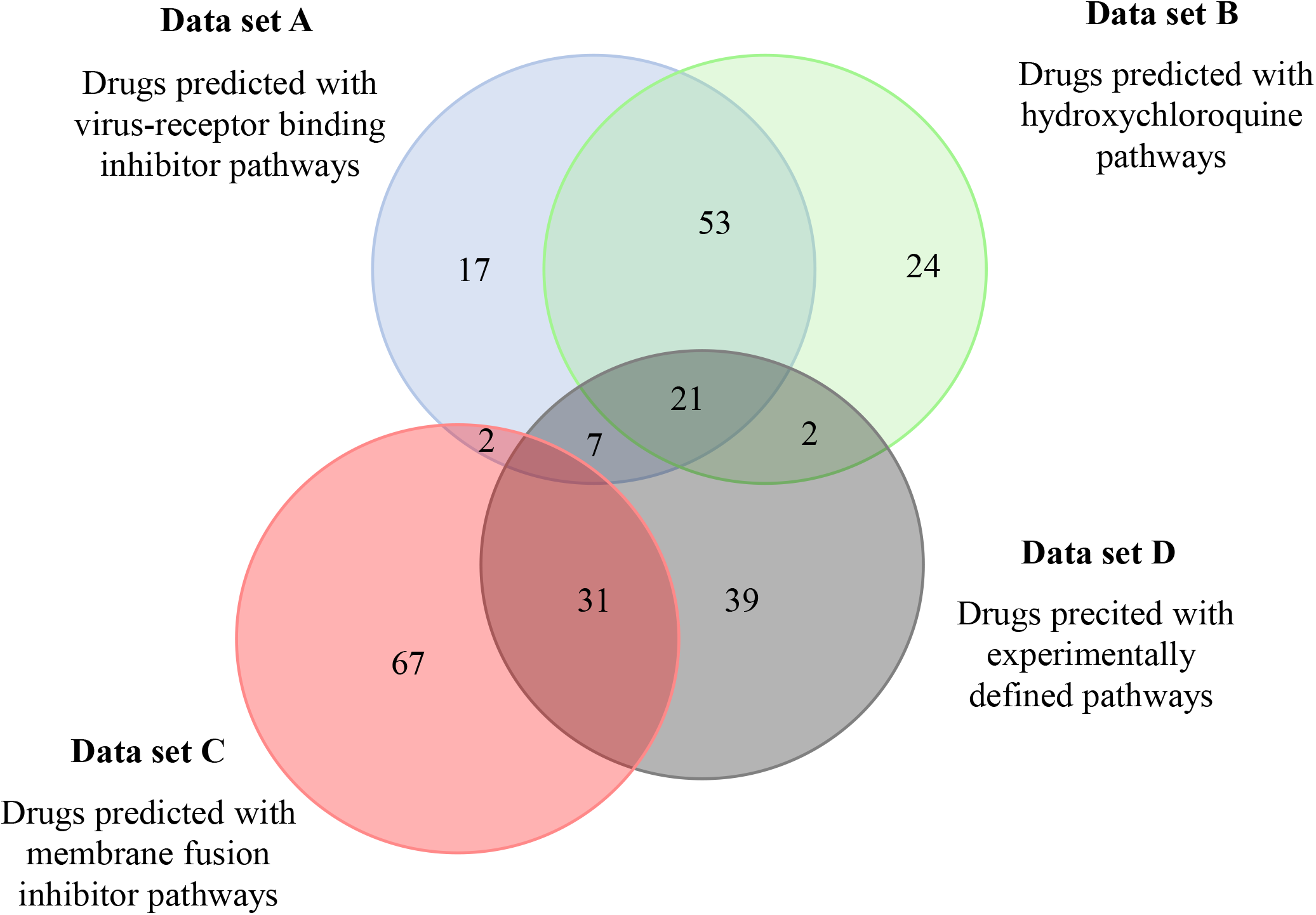
Overlap among drugs identified by the different pathway sets predicted to be involved in SARS-CoV-2 infection. Pathways relevant for SARS-CoV-2 infection predicted to be affected by three selected drugs were used to identify drugs from the DrugBank. Predicted drugs involved in this analysis are shown in Supplementary Material 3.

Anti-histamines were among the most prevalent drugs identified, both H1 and H2 receptor blockers. The H1-blocker anti-allergy medicine, azelastine, was predicted in three independent screens (Supplementary Material 3). Since azelastine has no major effect on normal physiology or concerning side-effect, it was in the focus of further studies.

We predicted genes to be involved in the action of azelastine and hydroxychloroquine and found that approximately two thirds of the genes obtained with azelastine overlapped with those identified with hydroxychloroquine, and were mainly related to immune response, which is in line with the anti-inflammatory effects of these two compounds, reported in the literature (Supplementary Material 4).^19,5^ However, we also found significant differences in the two gene sets that is indicative of distinct cellular mechanisms that may lead to antiviral effects (Supplementary Material 4).

### Anti-SARS-CoV-2 activity of azelastine in Vero E6 cell line

Azelastine was first tested for anti-viral effect in a gold standard assay of SARS-CoV-2 infection using the ACE2 expressing Vero E6 (African green monkey kidney epithelial) cells in the 0·4 and 25 μM concentration range, either with co-administration with the virus or as treatment after viral infection. Based on a semiquantitative assessment by microscopic examination of cells 48 hours post-infection, azelastine was effective in reducing the cytopathic effect in the 3 to 25 μM concentration range (Supplementary Fig. 1). The EC_50_ values of the anti-viral effects based on virus quantification from the culture supernatant were approximately 6 μM for both the co-administration and treatment settings (Fig. 4 A&B).

**Figure 4.**
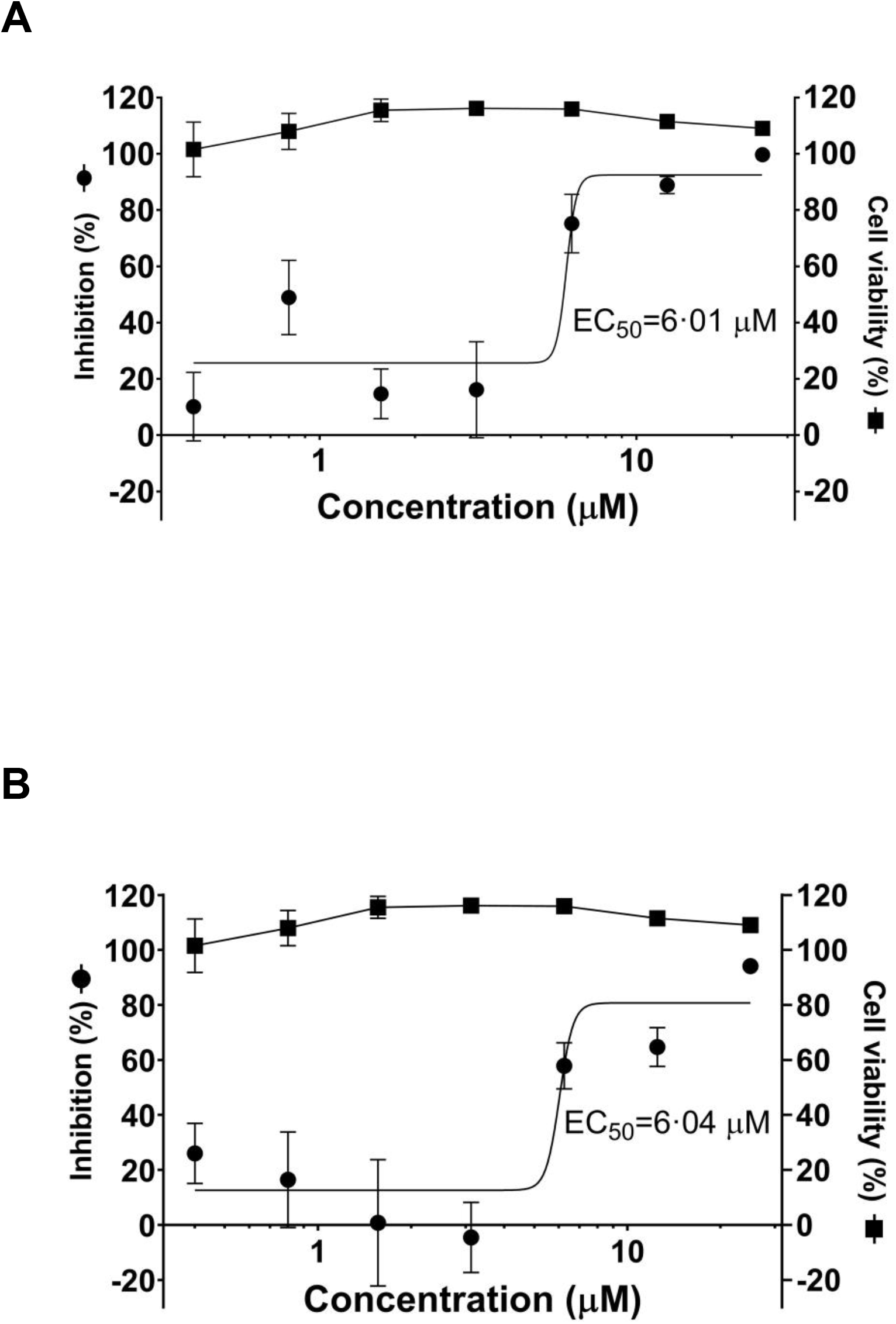
Azelastine is effective against SARS-CoV-2 infection in Vero E6 cell line. **A:** Vero E6 cells were infected with SARS-CoV-2 simultaneously with the addition of 0.4 to 25 μM of azelastine and after 30 min continued to be cultured without the virus in the presence of the respective concentrations of the drug. **B:** Cells were infected with SARS-CoV-2 for 30 min, and then virus was removed by washing and culture medium was exchanged to medium containing 0·4 to 25 μM of azelastine. After 48 hours, RNA was extracted from the the culture supernatant and was quantified by droplet digital PCR analysis. Cell viability was determined in control cultures without viral infection treated with azelastine. Graphs show mean±SEM from 5 biological replicates and 2 technical repeats from selected samples. EC_50_ values were calculated with the GraphPad Prism 8.4.3 software.

### Anti-SARS-CoV-2 activity of azelastine containing nasal spray using reconstituted human nasal tissue

Since azelastine is widely used in anti-allergy nasal sprays, we tested one of the commercially available products on reconstituted human 3D nasal tissue (MucilAir™). The tissue samples were first infected with the virus and then treated with five-fold diluted 0·1% nasal spray solution for 20 min in every 24 hours for three days. Microscopic analysis of tissues at 48 and 72 hours revealed reduced mucin production in infected cells relative to control cells (no virus or drug treatment) that was preserved in the presence of azelastine (Fig. 5). No difference in tissue morphology was detected in control and azelastine-treated cells (without virus infection) and cilia movement was also detected (data not shown).

**Figure 5.**
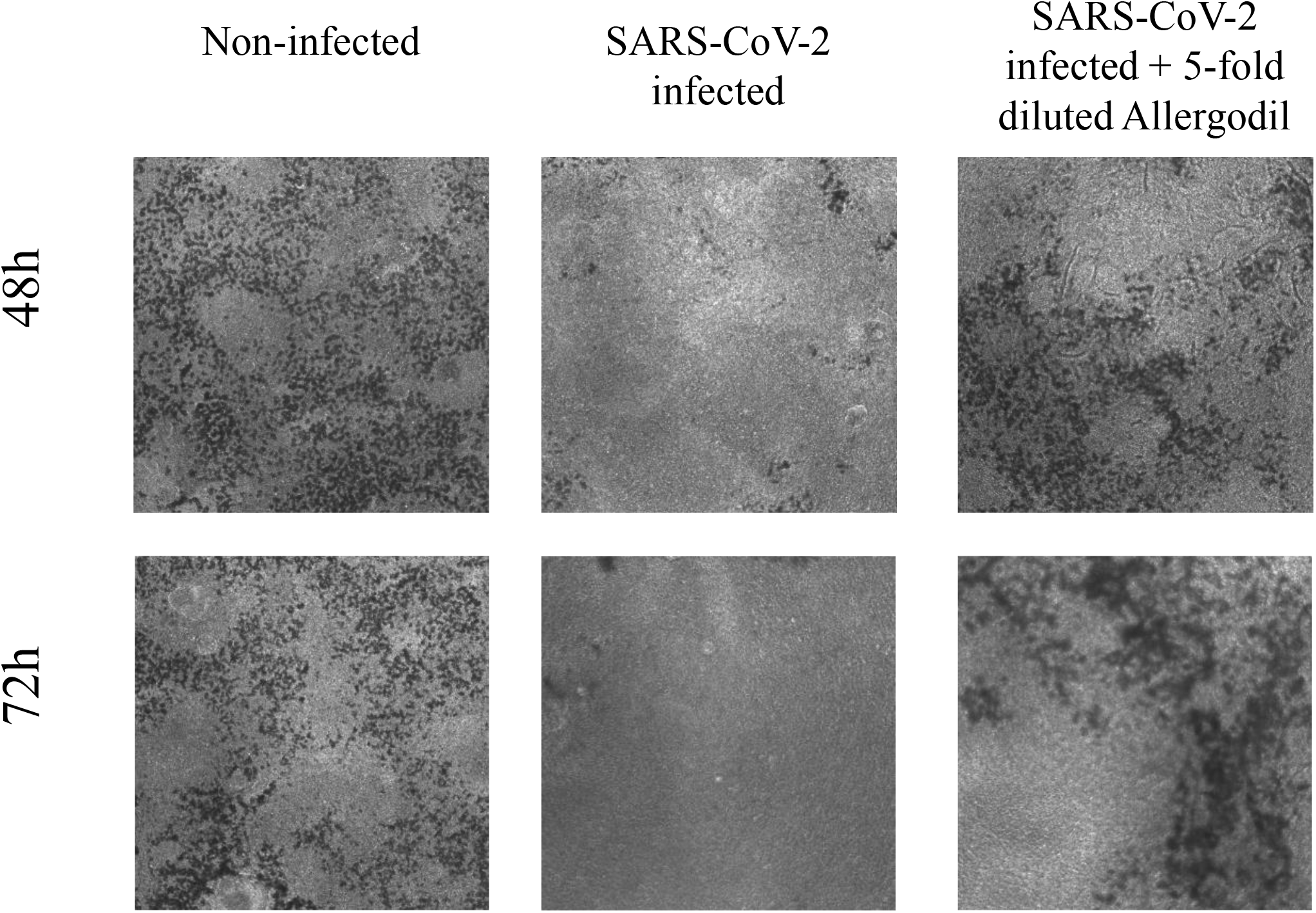
Azelastine blocks viral replication in SARS-CoV-2 infected reconstituted human nasal tissue. MucilAir™ human nasal tissue was infected with SARS-CoV-2 and subsequently treated with five-fold diluted 0·1% Azelastine-containing nasal spray solution for 20 min in every 24 hours for three days. Low resolution microscopic images of cultures after 48 and 72 hours of treatment.

Droplet Digital PCR analysis confirmed an effective SARS-CoV-2 infection and fast viral replication, reaching several thousand copies per microliter by 72 hours post-infection in the apical compartment of the tissue inserts (Table 1). Daily 20-minute treatment with azelastine (0·02%) drastically reduced the viral particle numbers (> 99·9% inhibition) by 48 and 72 hours post infection (Table 1).

**Table 1.** Viral RNA copy numbers in untreated and azelastine-treated nasal tissues.

| | Viral RNA copy/μl | | |
| --- | --- | --- | --- |
|  | 24 hpi | 48 hpi | 72 hpi |
| **Untreated** | 0.68 | 444.67 | 3521.33 |
| **Allergodil**<br>**five-fold diluted** | 0.053<br>(7.88%) | 0.027<br>(0.01%) | 0.05<br>(<0.01%) |

Values represent the average of triplicate samples. Results with azelastine treatment also expressed relative to the untreated infected control (%).

## Discussion

Selected by computational prediction and confirmed by *in vitro* experimental testing, we identified azelastine, an anti-allergy compound, broadly available in nasal formulation as a potential anti-COVID-19 remedy.

We pursued a pathway-centric drug repurposing approach to predict drugs with potential anti-COVID-19 activity. In our poly-pharmacological concept, a drug’s mode of action is based on the distribution of protein targets and the biochemical pathways they are part of. We argue this to be a more promising approach compared to conventional single target drug design strategies, particularly in view of the multi-factorial disease phenotype of COVID-19.

We screened the DrugBank database with three small molecules with proven SARS-CoV or SARS-CoV-2 activity to identify drugs with similar Shannon entropy vectors, and then used the information on the pathways these drugs are predicted to activate. The greatest overlap was seen with pathways known to be involved in bacterial, parasitic and viral infections and cellular processes involved in infectious diseases. Using the three predicted pathway profiles and one experimental ^16^, we identified 263 drugs, with 117 appearing in more than one screens. The major drug classes were anti-infectives (antivirals, antibiotics, antifungals), anti-histamines, anti-inflammatory drugs (non-steroids compounds), anti-hypertensive drugs, and CNS-targeting drugs, mainly anti-psychotics.

Interestingly, we found some overlap with drugs predicted by a complex network approach reported by the Barabási group, which relies on information about human protein binding partners where potential drug candidates are likely to perturb the interactome network relevant for viral infection ^15^. Although our methodology considers biochemical pathways independent of individual protein interaction events, 15 of the 81 top scoring and prioritized drugs from that study were also identified in our screens.

Several of the predicted drugs from this study are being investigated for efficacy in clinical studies. Most notably, dexamethasone has been proven to reduce mortality of COVID-19 patients on mechanical ventilation by one third.^2^ Two additional drugs predicted here, famotidine, an H2-blocker anti-histamine used to decrease gastric acid production and telmisartan, an angiotensin receptor blocker anti-hypertensive drug, have been recently reported to improve morbidity in hospitalized patients infected with SARS-CoV-2.^21,22^ The synthetic serine protease inhibitor nafamostat mesylate, approved for pancreatitis and also used as a fast-acting anticoagulant, is being tested for efficacy against COVID-19.^23^ Its anti-SARS-CoV-2 activity is explained by the inhibition of the plasma membrane-associated serine protease, TMPRSS2, that is employed by the virus to cleave and activate the Spike protein required for virus entry in lung epithelial cells.^23^ Nafamostat proved to be more efficacious *in vitro* than another serine protease inhibitor, camostat, also being tested clinically.^23^

Several of the drugs predicted in this study have been shown to have activity against coronaviruses *in vitro*. Cepharanthine, a natural compound with anti-inflammatory and anti-neoplastic activities was identified in a recent large-scale high throughput COVID-19 drug repurposing project using libraries of ∼3000 drugs (∼1000 of these FDA approved drug) ^8^ and also by others.^24,25^ Previous studies demonstrated its inhibitory effects on both SARS-CoV ^26^ and HCoV-OC43.^27^ Clofazimine, an antibiotic used against leprosy was identified as one of the 21 active drugs against SARS-CoV-2 (that reduced viral replication by at least 40% and showed dose-response) selected in a large-scale screening effort with 12,000 clinical stage or FDA-approved small molecules.^14^

From the predicted drugs with unproven anti-SARS-CoV-2 activity, we focused on azelastine (testing of other drugs is ongoing). This choice was driven by the following: 1., favorable side-effect profile, 2^nd^ generation non-sedating in most individuals, 2., no major effect on normal physiology, since the indication is to alleviate allergy symptoms, 3., broad availability, wide use, low cost; 4., availability in a nasal formulation for potential effect on nasal colonization; 5., several H1 and H2 blockers identified in our screens and others, e.g. ebastine ^8,28^; and 6., positive clinical data with the H2 receptor antagonist famotidine.^21^

Here we provide experimental proof that azelastine is effective against SARS-CoV-2 infection in the most widely employed *in vitro* assay system with Vero E6 cells with comparable EC_50_ value (∼6 μM) determined for chloroquine, lopinavir, and remdesivir (7 to 11 μM) using the same cell line.^25^ The great disappointment with chloroquine/hydroxychloroquine in clinical efficacy studies raises the concern about the predictive value of such *in vitro* results. Recent data revealed that the choice of cells in SARS-CoV-2 infection assay has great influence on the outcome of drug repurposing testing and showed that hydroxychloroquine was not active against the virus in human respiratory epithelial cells.^7,8^ This data suggests that inhibition of endosomal acidification, the mode of action of hydroxychloroquine is not relevant to respiratory epithelial cells.

Based on the GO term analysis and the predicted genes involved, azelastine is likely to exploit different biological processes compared to hydroxychloroquine. The exact molecular mechanism of its anti-viral effect is currently not yet delineated and needs further scientific exploration.

Importantly, we confirmed the efficacy of azelastine in human respiratory epithelial cells using a highly relevant *in vitro* model, the reconstituted human nasal tissue. In this model we simulated the clinical situation of nasal colonization by SARS-CoV-2 and observed the complete halting of viral propagation even after the first treatment for 20 min with a five-fold diluted commercial azelastine-containing nasal spray solution.

Azelastine is a multifaceted drug. It is best known as a histamine H1 receptor blocker, acting not as an antagonist but as inverse agonist, decreasing H1 receptor constitutive activity.^19^ However, azelastine has also general anti-inflammatory effects, mainly exerted via mast cell stabilization and inhibition of leukotriene and pro-inflammatory cytokine production.^19^ Importantly, mast cells are the main sources of cytokine release that leads to lung damage in SARS-CoV-2 and it has been speculated that mast cell stabilisers may also attenuate pulmonary complications, fatal inflammation and death in COVID-19.^29^ Therefore, azelastine’s potential beneficial effects in COVID-19 are expected to be the combination of antiviral and host-mediated actions. Recent clinical data with the H2 receptor antagonist famotidine, supports the potential therapeutic benefit of anti-histamines in COVID-19 patients.^21^

The synergy between H1 blockers and glucocorticoids due to the interaction between H1 receptor- and glucocorticoid receptor-activated pathways, is well recognized clinically, and the two types of drugs are often used in combination.^30^ Given the clinical effectiveness of dexamethasone on COVID-19 morbidity and mortality, testing of anti-histamines and anti-inflammatory steroids in combination is highly warranted.

The major implication of our findings is that a widely available nasal spray formulation containing azelastine might be an immediate solution to prevent and treat nasal colonization with SARS-CoV-2 therefore may have a great impact on the viral spread within the affected person (nose to lung) as well as within the population. This potential needs to be confirmed in clinical studies.

## Acknowledgement

We thank Katalin Gombos MD and Rita Csepregi (Dept of Laboratory Medicine, Faculty of Medicine, University of Pécs) for their kind technical support.

## Funding sources

This research was funded by CEBINA GmbH and the Hungarian Scientific Research Fund OTKA KH129599, European Social Fund: Comprehensive Development for Implementing Smart Specialization Strategies at the University of Pécs (EFOP-3.6.1.-16-2016-00004) andthe Higher Education Institutional Excellence Program of the Ministry for Innovation and Technology in Hungary, within the framework of the “Innovation for a sustainable life and environment” thematic program of the University of Pécs (TUDFO/47138/2019-ITM).

---

**Figure S1.**
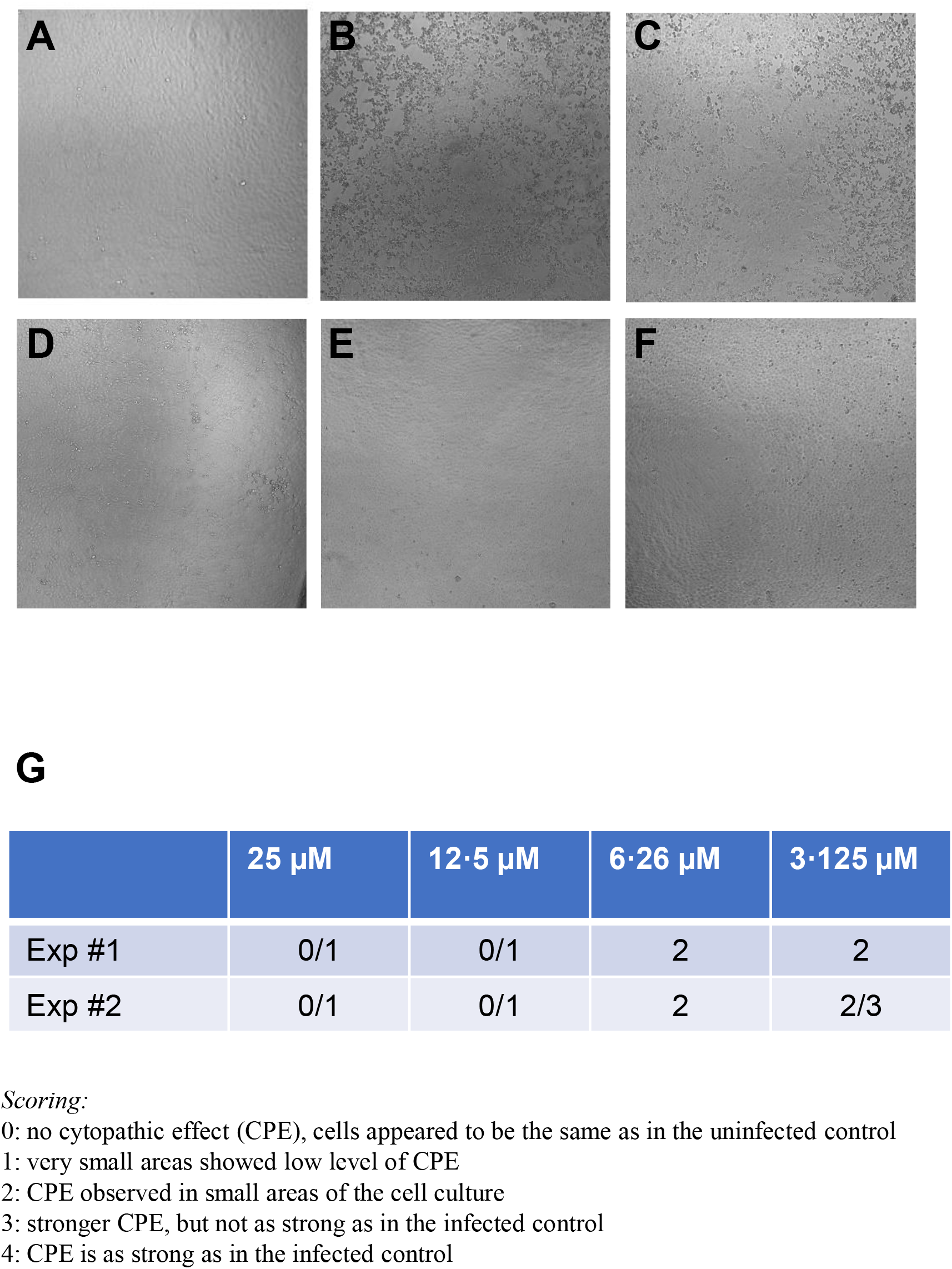
Prevention of SARS-CoV-2 induced cytopathic effect by azelastine. Vero E6 cells were infected with SARS-CoV-2 simultaneously with the addition of 0·4 to 25 μM of azelastine and continued to be cultured without the virus in the presence of the respective concentrations of the drugs. Cytopathic effect was assessed by light field microscopic examination of cultures 48 hours post-infection. **A:** uninfected (negative) control, **B:** virus infected (positive) control, **C:** virus + 3·125 μM azelastine, **D:** virus + 6·25 μM azelastine, **E:** virus + 12·5 μM azelastine, **F:** virus + 25 μM azelastine. **G:** Scoring system and summary of cytopathic effect in the presence of azelastine (results of two independent experiments)

## Notes

### Competing Interest Statement

EN, GN and VS are employees and shareholders of CEBINA GmbH. TG is employee and shareholder of Calyxha GmbH. RK receives consultancy fee from Calyxha GmbH and holds shares in the company. This research was funded by CEBINA GmbH, the Hungarian Scientific Research Fund OTKA KH129599, the European Social Fund: Comprehensive Development for Implementing Smart Specialization Strategies at the University of Pecs (EFOP-3.6.1.-16-2016-00004) and the Higher Education Institutional Excellence Program of the Ministry for Innovation and Technology in Hungary, within the framework of the "Innovation for a sustainable life and environment" thematic program of the University of Pecs (TUDFO/47138/2019-ITM).

