## Supplementary Material 1 for "The Anti-histamine Azelastine, Identified by Computational Drug Repurposing, Inhibits SARS-CoV-2 Infection in Reconstituted Human Nasal Tissue In Vitro"

**Computational method to predict drug pathway profiles**

The approach was based on the Shannon-Entropy Descriptor (SHED) concept. In the conventional SHED approach^1^ the chemical structure is converted into a 2D topological graph in which the nodes correspond to the atoms in the drug and edges connecting two nodes indicate the existence of a chemical bond. From this graph the shortest path length between each and every residue pair (characterized by their atom centered features) is calculated and stored as a function of feature pair. The Shannon-entropy quantifies the variability of feature-pair distributions in the molecule. Here, we have considerably extended the list of atom-centered features: (Lipophylic (L), positively charged (P), negatively charged (N), Hydrogen-Bond Donor (D), Hydrogen-Bond Acceptor (A) and simultaneous Hydrogen-Bond Donor/Acceptor (AD). A given atom is thus described via a 6-letter string consisting of (0,1), where 1 indicates the presence of a given feature. For example, a sp3-carbon atom is given by (100000), while a carboxylic oxygen reads as (001010) indicating the negative charge and the hydrogen bond acceptor capability. In total 25 feature pairs are used. Drug information is stored as a 1D Shannon entropy vector and thus allows for large-scale applications. Pairwise drug similarities were quantified by calculating the Euclidean distance. To obtain the biochemical pathway profile of a query molecule we screened the DrugBank database^2^, a repository of approved drugs and their experimentally verified protein targets. In a first step drug analogs from the DrugBank database are identified using the Shannon entropy vector (euclidean similarity cutoff: 0.25). Next, the experimentally verified protein targets for the identified DrugBank analogs are used to derive information about the involved KEGG biochemical pathways. KEGG pathways are quantified by enumerating how often they are found in the list of identified DrugBank analogs. The numbers are normalized to the number of the most prevalent KEGG pathway (typically metabolic pathway or neuroactive ligand-receptor interaction pathway). Finally, the normalized pathway numbers are referenced (difference) to the statistical abundance of KEGG pathway obtained for the entire DrugBank dataset. The statistical abundance of the KEGG pathways was obtained by predicting the KEGG pathway profile (as described above) for all of the drugs in the DrugBank database.

The resulting ranked profile signatures were then used to score corresponding genes of azelastine-HCL and hydroxychloroquine. These genes were identified and described for homo sapiens using the HGNC Database, HUGO Gene Nomenclature Committee (HGNC) of the European Bioinformatics Institute (EMBL-EBI) in the biomaRt package.^3^ For azelastine-HCL 201 and for hydroxychloroquine 185 genes were identified with an overlap of 129 genes. For azelastine-HCL, 72 genes were exclusively identified and 56 genes were exclusive for hydroxychloroquine (Supplementary Material 4).

**References: Computational method**

1	Gregori-Puigjané E, Mestres J. SHED:  Shannon Entropy Descriptors from Topological Feature Distributions. *J Chem Inf Model* 2006; **46**: 1615–22.

2	Wishart DS, Feunang YD, Guo AC, *et al.* DrugBank 5.0: a major update to the DrugBank database for 2018. *Nucleic Acids Res* 2017; **46**: gkx1037-.

3	Durinck S, Spellman PT, Birney E, Huber W. Mapping identifiers for the integration of genomic datasets with the R/Bioconductor package biomaRt. *Nat Protoc* 2009; **4**: 1184–91.
